# Disambiguation of two-tone images aligns object representations in the EEG

**DOI:** 10.64898/2026.06.05.730193

**Authors:** Roman Kessler, Johanna J. Finnemann, Michael A. Skeide

## Abstract

Electrophysiological responses to visual objects carry information about stimulus identity and semantic category, but it remains difficult to know whether such information represents conceptual knowledge or merely regularities in physical image features. Here, we presented two-tone images before and after disambiguation via corresponding photographs to seventeen healthy participants while recording EEG to dissociate the learned image concept from its physical properties. After disambiguation, EEG responses to learned two-tone images became more similar to the responses oftheir photo counterparts. This effect emerged approximately 100 ms after stimulus onset and peaked at 350 ms. However, we found no evidence for a generalization of the response pattern to the stimulus semantic category. Furthermore, alignment to the photo counterparts was only detectable after explicit visual disambiguation rather than spontaneous recognition. Together, these findings indicate that EEG can capture specific learned conceptual modulations of responses to visually degraded stimuli, while also showing that low-level visual properties dominate the representational structure.

## Introduction

Patterns of brain activity carry information about the semantic content of visual stimuli ^1–8^. However, when neural activity is used to distinguish between visual stimuli it is often unclear whether the signals carry information about semantic content or merely variation in low-level visual features ^9^. As a consequence, studying category discrimination on a neural level requires a method to separate the semantics of the object (e.g., the category) from the remaining image properties, i.e., the low-level image statistics. This is particularly challenging, as physically harmonizing all low-level image features of visual stimuli inevitably incorporates equating the entire images. For example, cat pictures will always encompass one or more low-level image statistics that are different from those of brutalist architecture.

The issue of identifying the semantic content of the neural response is particularly relevant for studies of image category perception in infants or clinical cohorts where behavioural confirmation of conceptual knowledge is challenging. By separating low-level image statistics from semantics one could assess on a neurophysiological level whether object categories can be distinguished at a particular age, or whether a learning intervention changes the semantic neural representation. Such a separation of low-level features and semantics could be achieved on an analytical level, for instance by including low-level image statistics as nuisance variables in the computation of the similarity metrics ^10,11^. Previous studies, however, only controlled a few out of many possible low-level features that constitute an image in its entirety. A different approach is to experimentally separate the image statistics from the concept in the paradigm design, for instance, by changing the concept associated with the same physical stimulus ^12^.

In the present study we followed this latter approach and leveraged two-tone (Mooney) images ^13^, whose interpretations shift from ambiguous to meaningful when observers are provided with supplementary information that reveals the underlying object, such as the corresponding photo image (i.e., template) or verbal instruction ^14–16^. Because the two-tone stimuli remain physically unchanged across the two knowledge states, they offer the opportunity to vary semantic interpretation without altering low-level visual features. The sudden shift from ambiguity to meaningful perception reflects the visual system’s hierarchical and recurrent architecture, in which high-level object knowledge can swiftly reshape the interpretation of impoverished sensory input. High-level object representations engaged during two-tone disambiguation have been shown to interact with early visual areas, indicating bidirectional influences between conceptual interpretation and sensory processing ^17^. Such interactions may dynamically weight the reliability of the sensory evidence ^18^ and sharpen perceptual representations according to contextual information ^19^.

Functional magnetic resonance imaging (fMRI) has revealed that neural responses to disambiguated two-tone images and corresponding photo images align in a widespread cortical network ^20,21^. For electroencephalographic (EEG) response profiles, this alignment is less clear. Although recognition is associated with power changes in different frequency bands ^16,22^, other EEG studies investigating the alignment of neural responses to two-tone images to their photo image counterparts showed small effects in their preliminary results and were not further pursued ^23^. Another study demonstrated such an alignment using magnetoencephalography (MEG) ^24^. However, EEG is often the only suitable method to investigate challenging cohorts such as infants or certain clinical patients. EEG also offers high temporal resolution that allows rapid presentations of stimuli, opening the opportunity to cover a larger stimulus space than previous studies ^21,23^. Notably, previous studies rely on slow event-related presentations, whereas the zeitgeist of object decoding assimilates rapid serial presentations ^25,26^, capitalizing on the parallel co-existence of several neural stimulus representations ^27^.

Establishing under which circumstances EEG response patterns reflect semantic knowledge such as stimulus identity or category instead of merely the low-level stimulus properties is a prerequisite for interpreting representational analyses. The present study therefore aims at demonstrating how manipulation of the semantic concept changes the electrophysiological representational structure. To this end, we presented two-tone images and their photo image counterparts to a group of healthy participants during EEG recording at a dense presentation rate of 2.5 Hz. Our primary hypothesis was that, after disambiguation, neural responses to two-tone images would become more similar to responses elicited by their corresponding photo images, indicating knowledge-dependent changes in stimulus representations, i.e., template alignment. In addition, we made two exploratory predictions concerning whether this representational alignment would generalize beyond the individual stimulus towards a common semantic category representation, and whether spontaneous recognition leads to a similar template alignment as explicit visual disambiguation.

Our results demonstrate an alignment of electrophysiological responses to two-tone images and their corresponding photographs after disambiguation. However, we could neither find evidence for the semantic category information being encoded in the electrophysiological signal after disambiguation nor an enhanced alignment between neural responses to spontaneously recognized two-tone images and their corresponding template images.

## Results

Seventeen participants completed an EEG experiment consisting of three phases. In the pre-disambiguation phase (PRE), participants viewed ten different two-tone images from five semantic categories (cow, croissant, hands, human, penguin; Fig. S1) presented using rapid serial visual presentation at 2.5 Hz (Fig. 1). In the template phase (TPL), participants viewed the corresponding grayscale photo images presented at the same rate. In the post-disambiguation phase (POST), participants were again exposed to the same two-tone images, but they had now learned the identity of each image by explicit visual transitions between the two image types (i.e., disambiguation). Disambiguation was performed prior to the TPL phase and before each POST block (Fig. 1). Participants provided verbal responses regarding the identity of the two-tone images after both the PRE and POST phases. These responses were used to assess recognition and to classify stimuli into different groups, i.e., (re-)learned, not (re-)learned, and spontaneously recognized (Table 1).

**Fig. 1.**
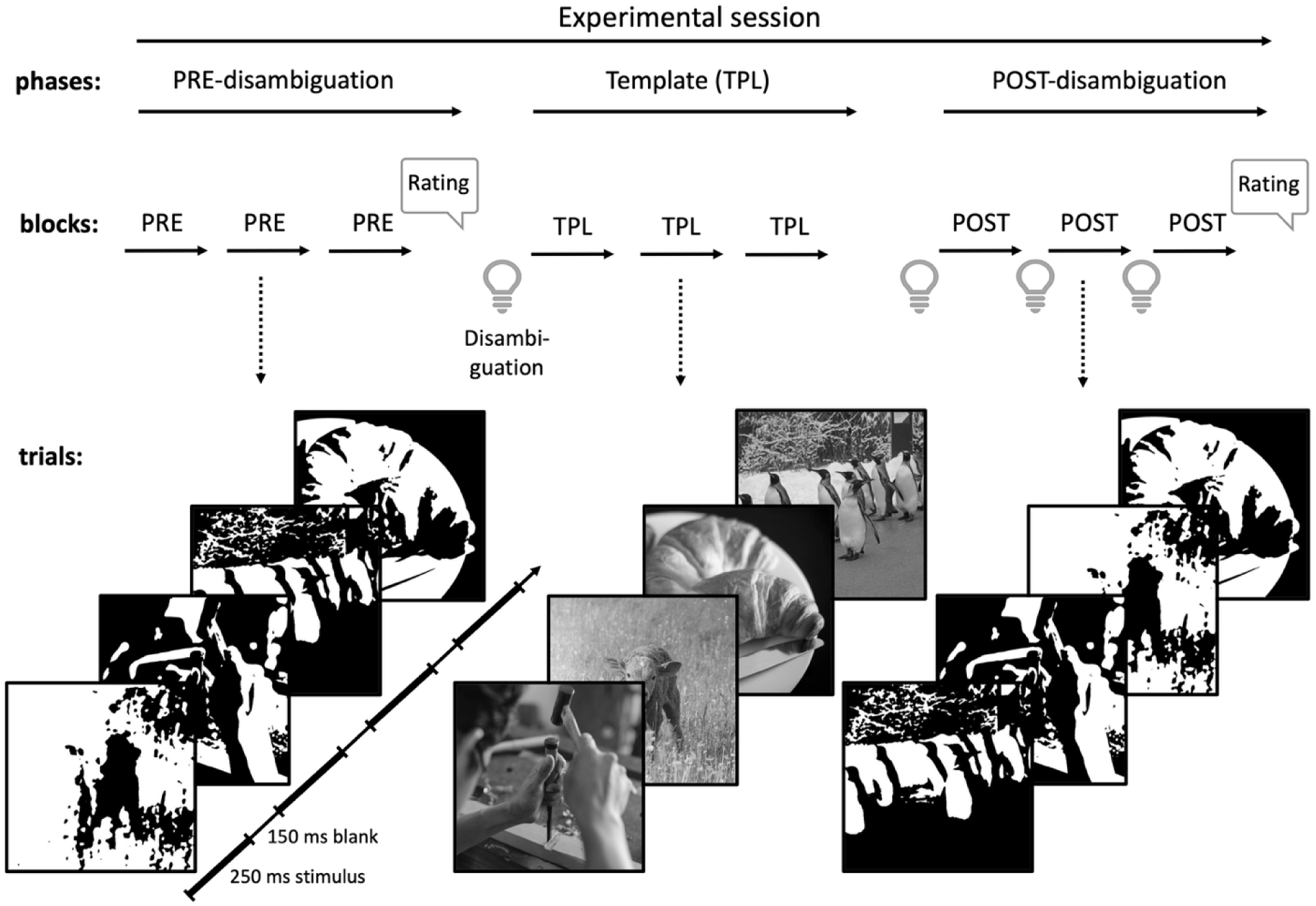
Experimental design. The experimental session was divided into three phases (upper panel), each comprising three experimental blocks (middle panel). Two-tone images were presented in the PRE- and POST-disambiguation phases, and their corresponding photo images were presented in the template (TPL) phase. Within each block, images were presented in random order at a rate of 2.5 Hz (250 ms stimulus, 150 ms inter-stimulus-interval, lower panel). Before the TPL phase and before each POST block, the two-tone images were disambiguated by a continuous fade between the two-tone image and its corresponding photo image (light bulbs). After PRE and POST phases, the participants were asked about stimulus identity (speech bubbles).

**Table 1.**
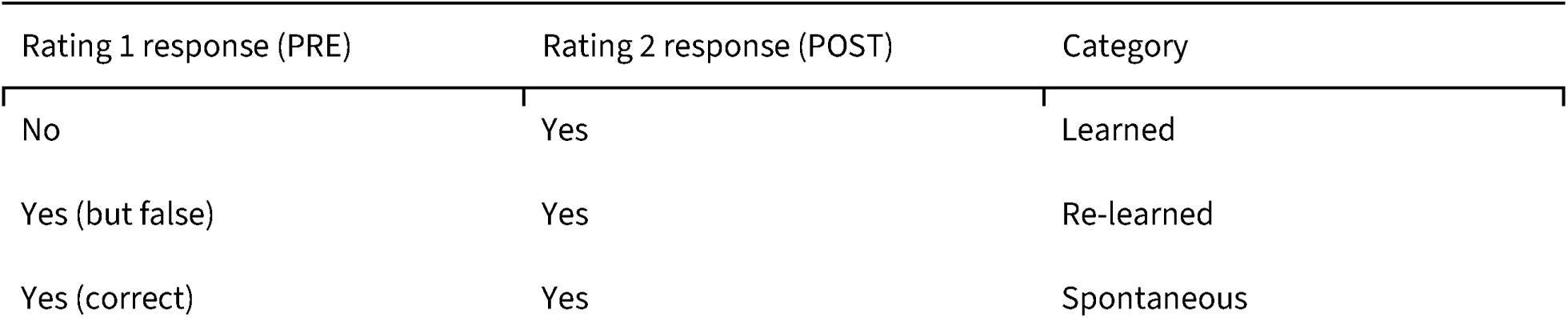
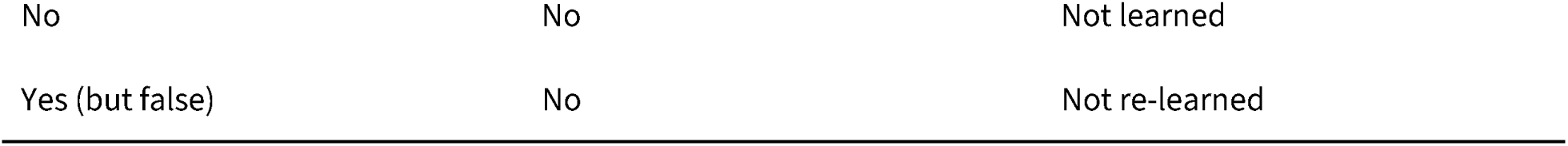
Categorization of responses to the stimuli. The responses given during the rating after the PRE and POST phases were used to categorize each stimulus for each participant. The questions were as follows: Rating 1 (PRE): “Did you recognize anything in this stimulus when it was shown during the previous runs? And if so, what did you see?” Rating 2 (POST): “Did you see the learned stimulus identity in the images within the previous runs?”. For confirmation, we asked for the image identities. No other combination of responses than the ones illustrated occurred.

| Rating 1 response (PRE) | Rating 2 response (POST) | Category |
|---|---|---|
| No | Yes | Learned |
| Yes (but false) | Yes | Re-learned |
| Yes (correct) | Yes | Spontaneous |
| No | No | Not learned |
| Yes (but false) | No | Not re-learned |

### Behavioral results

All participants completed the entire experiment, but varied regarding which and how many stimuli they spontaneously recognized before disambiguation, which they successfully learned or re-learned after disambiguation, or which they did not learn. Table 1 illustrates how participants’ responses were categorized. Participants spontaneously recognized 0 to 3 out of 10 stimuli, were not able to learn 0 to 1 out of 10 stimuli, and learned or re-learned the remaining 6 to 10 out of 10 stimuli. Figure 2B illustrates how each stimulus was categorized for each participant (vertical panels). Table S1 further summarizes how many participants did or did not (re-)learn, or spontaneously recognized each stimulus.

**Fig. 2.**
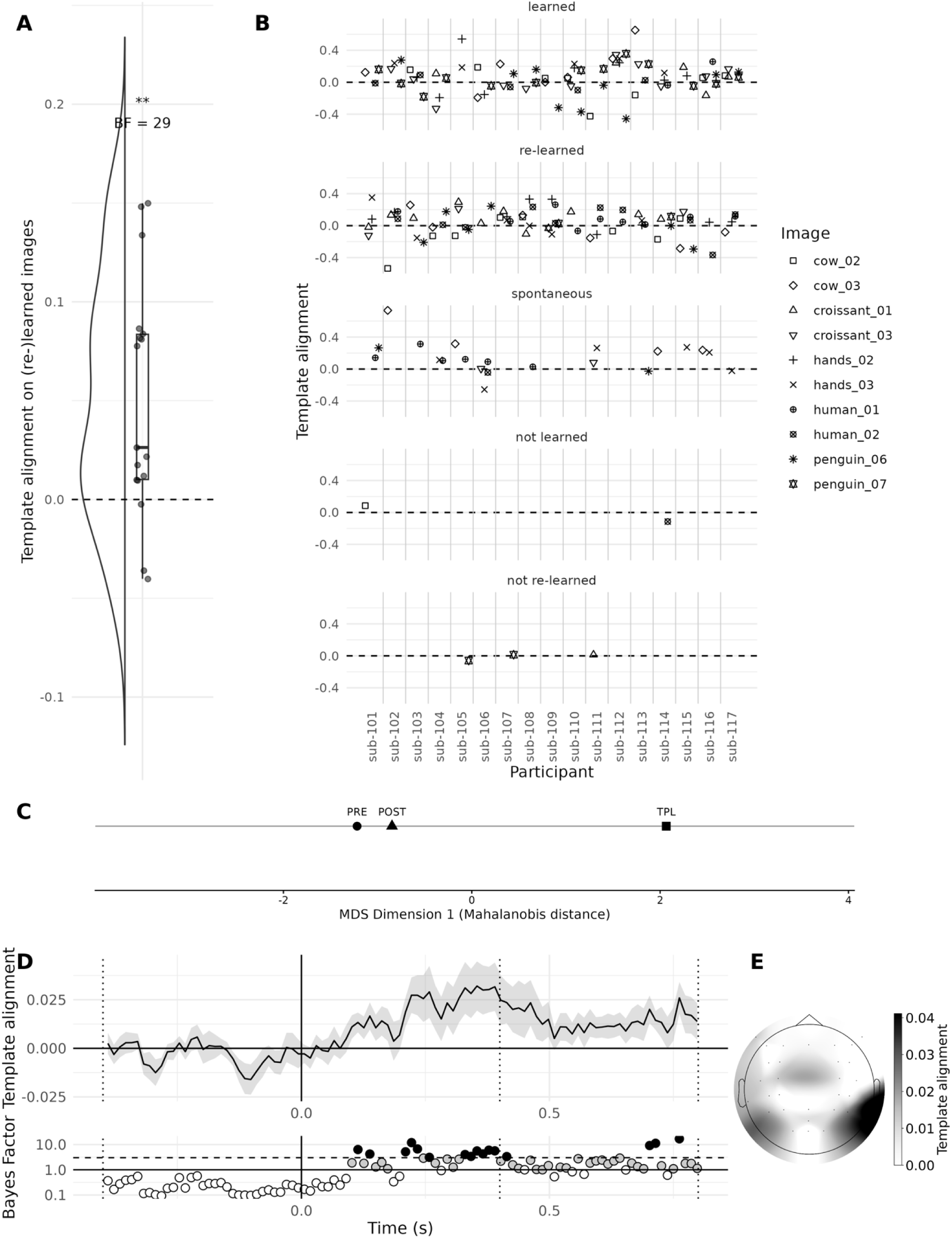
Template alignment. **A**: Participant-wise template alignment of (re-)learned stimuli. The y axis illustrates the difference (PRE – POST) in neural response dissimilarity to the template image, i.e., template alignment in the window 0.1 – 0.4 s post-stimulus onset, averaged across each participant’s learned & re-learned images. Boxplot corpus illustrates 0.25, 0.5, and 0.75 quantiles. Whiskers span all data points within 1.5 times the inter-quantile-range. BF = Bayes Factor. Asterisks indicate significance levels (** p < 0.01, one-tailed). **B:** Stimulus- and participant-wise template alignment values. Vertical panels illustrate the different stimulus categories based on participant ratings (Table 1). The x axes depict the different participants. The y axes illustrate the difference (PRE – POST) in neural response dissimilarity to the template image, i.e., template alignment. Shapes illustrate the different stimulus images. **C:** Estimated multidimensional scaling of distance values of learned and re-learned stimuli (with only one dimension). D & E: Temporal trajectory and spatial topography of template alignment. **D:** The upper panel shows the mean template alignment value (y axis) per time point (x axis) calculated across all electrodes with standard errors across participants. The lower panel shows the respective BFs (y axis, logarithmic scale, one-tailed), with grey circles indicating a BF > 1 and black circles indicating a BF > 3. The horizontal solid line at y = 1 indicates the points where the likelihood for a positive template alignment value is equal to the likelihood of a null or negative template alignment value, whereas the horizontal dashed line indicates a BF = 3. The solid vertical line indicates the onset of the stimulus of interest. Dotted vertical lines indicate the onsets of (randomized) preceding and succeeding stimuli. E: Mean template alignment values were computed for each electrode individually and across the entire time window of interest (0.1 – 0.4 s). No electrode exceeded a BF of 3.

### Template alignment

We analyzed how the neural responses to all stimuli differed across experimental phases for each participant, with a focus on learned and re-learned stimuli. First, we computed the Mahalanobis distance of neural responses between POST and TPL, and subtracted it from the Mahalanobis distance between PRE and TPL, all within a post-stimulus onset time-window of 0.1 – 0.4 s. This difference measure quantifies what we label template alignment (eq. 1a, cf. Methods). The analysis window starts shortly after stimulus onset, when initial neural responses to that stimulus can be detected. The window ends with the onset of the next stimulus and thus before the signal might be influenced by the neural response to the next stimulus. Thus the window covers an interval in which both perceptual and conceptual processing can be delineated from the electrophysiological signal ^10,28^. The calculation includes subtracting stimulus-unspecific, phase-dependent components via cross-stimulus pairings (eq. 1b, cf. Methods). A positive template alignment value indicates that the neural response to the two-tone image became more similar to the neural response to the respective template image. All analyses in the present study were conducted separately for each participant, as individual differences between neural representations were expected. The participant-wise results were passed to the group level for statistical analysis. Pair-wise statistical analyses were always one-tailed in the direction of the respective hypothesis.

Figure 2A illustrates the average template alignment value of each participant, including all learned and re-learned stimuli per participant. We obtained a Bayes Factor (BF) of 29.0, indicating strong evidence that the true value is positive. A one-sided Wilcoxon signed-rank test confirmed that template alignment was significantly greater than zero (V = 135, p = 0.0019). The observed effect size was g = 0.80 with a 95% confidence interval (CI) of [0.27, 1.33]. Figure 2B illustrates the template alignment values for each stimulus and participant. Vertical panels depict how each stimulus was classified according to Table 1 for each participant. (Re-)learned stimuli revealed positive template alignment values. A sequential trial accumulation analysis revealed an increase of this effect with the number of stimulus repetitions, with about 25% of currently used 240 repetitions leading to a BF of 3, and around 70% to a BF of 10 (Fig. S2). A sensitivity analysis on the experimental design indicates that it is well-suited to detect effect sizes of 0.71 or larger with probability of > 80% (Fig. S3). Furthermore, converging results were obtained when using different distance metrics, or when limiting the analyses to posterior electrodes (see Supplementary Information).

Our next step was to compare Mahalanobis distance values across the experimental phases, by contrasting electrophysiological distance values corresponding to PRE vs. TPL, POST vs. TPL, and PRE vs. POST. Figure S4 illustrates all raw distance values from all stimuli. Distances between the PRE and POST electrophysiological responses were consistently lower than the distances between responses to PRE and TPL, or POST and TPL. As the PRE and POST stimuli differ only in their semantic interpretation (and their position in the experimental sequence), whereas the other comparisons also differ in their visual features, this result illustrates that the low-level image statistics explain a larger proportion of variance in the electrophysiological signal than the concept shared by POST and TPL ^23^. We used the distance values of all learned and re-learned stimuli across participants to estimate a multidimensional scaling model with only one dimension. Figure 2C illustrates that PRE and POST stimuli are situated in vicinity to each other, whereas TPL stimuli are situated further away. POST stimuli are situated slightly closer to the TPL stimuli than the PRE stimuli are. The distance from POST to TPL was about 8 times the distance from PRE to POST stimuli, indicating that disambiguation only slightly alters the electrophysiological response to two-tone images. This pattern was observed in 15 out of 17 individual participants (Fig. S5).

To explore the temporal trajectory of template alignment, we applied the same formula (eqs. 1a&b) for consecutive 12–millisecond time-windows of the experimental trials ^29^. Figure 2D illustrates the mean time-point-wise template alignment across participants and their corresponding BFs, spanning 3 stimulus cycles (-0.4 to 0.8 s) around stimulus onset. The mean template alignment value across participants peaks between 300 and 400 ms after stimulus onset, and remains positive until the end of the investigated time window. Evidence for template alignment surpasses the evidence of the alternative (i.e., template alignment being zero or negative) shortly after stimulus onset. The BF fluctuates in a range between roughly 1 and 16 until 800 ms after stimulus onset, indicating traces of template alignment persisting even beyond the next stimulus onset, in line with the notion of concurrent representational existence of several stimuli as in rapid serial visual presentation paradigms ^27^.

To better understand possible spatial origins of template alignment responses, we computed electrode-wise template alignment values, using the entire time-window of interest (0.1 – 0.4 s). Positive template alignment values were observed on posterior lateral channels, but none exceeded a BF of 3 (Fig. 2E). We repeated this analysis within 100 ms consecutive time windows, illustrating that the most positive template alignment values originated from distributed, mostly posterior lateral electrodes in an extended time-window starting shortly after stimulus onset (Fig. S6).

### Category selectivity

As a follow-up question we aspired to answer whether the alignment generalized beyond the exact stimulus, i.e., if disambiguation not only leads to stimulus-level representational encoding, but also triggers a semantic category-level encoding. As category selectivity should only emerge in learned two-tone images, we only focused on the (re-)learned stimuli. Specifically, we determined if the neural responses of two different (re-)learned two-tone stimuli belonging to the same category (e.g., penguin_06 and penguin_07, Table S1) were more similar (less dissimilar) to each other, than to neural responses to stimuli of other categories. Category selectivity was computed separately for each experimental phase and participant (eq. 2, cf. Methods).

Figure 3A illustrates the category selectivity values for each participant and experimental phase. Experimental phases differed in category selectivity (Friedman-Test: ^2^(2) = 15.6, p = 0.0004). Category selectivity was not significantly higher in the POST than in the PRE phase (V = 50, p = 0.897), with an effect size of g = -0.29 (95% CI = [-0.75, 0.18]). Bayesian analysis also revealed evidence against an increase in category selectivity via disambiguation (BF = 0.1238). As expected, the TPL phase exhibited higher category selectivity than the PRE phase (V = 142, p = 0.0006, g = 0.76 95% CI = [0.23, 1.28], BF = 20.9), and the POST phase (V = 148, p = 0.0002, g = 0.85 95% CI = [0.30, 1.38], BF = 42.4). P values were false-discovery-rate adjusted using Benjamini-Hochberg correction ^30^. To investigate if perceptual similarity systematically biased category selectivity values (i.e., y axis offsets in Figure 3A), we compared pixel-wise correlation between the stimuli within and between categories, and between photos and two-tone images. Pixel-wise similarity did not differ between within- and between-category pairs (photos: p = 1.00; two-tone: p = 1.00, Holm-corrected), between image types (p = 0.47), nor was there an interaction (p = 0.62, Fig. S7).

**Fig. 3.**
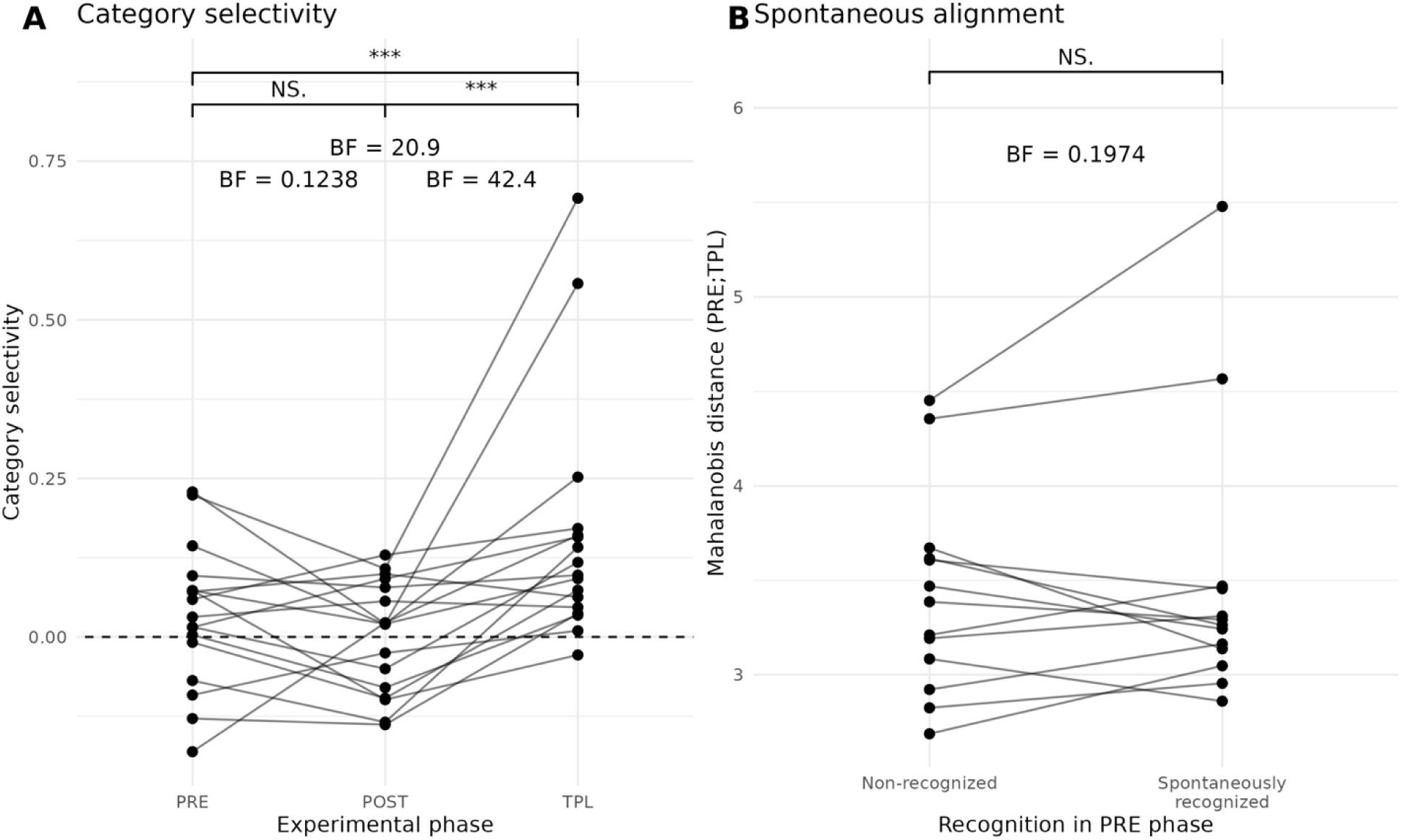
Category selectivity and spontaneous alignment. **A:** Category selectivity. The x axis illustrates the different experimental phases (PRE-disambiguation, POST-disambiguation, TPL: Template). The y axis illustrates the category selectivity (between-category minus within-category distances) for each participant. **B:** Spontaneous alignment. The x axis shows the two groups of stimuli, either those not recognized or those spontaneously recognized in the PRE phase. The y axis illustrates the participant-wise average dissimilarity of neural responses to those of their respective template images. BF = Bayes Factor. Asterisks indicate levels of significance (*** p < 0.001, one-tailed; NS. not significant)

### Spontaneous alignment

The final step was to explore potential actuators of template alignment, i.e., if neural representations align only after exposure to the template image or whether it evolves spontaneously when recognized. In particular, we tested whether the neural responses to recognized stimuli in the PRE phase were more similar to the neural responses of the respective photographs than those of their non-recognized counterparts. Accordingly, we calculated the mean dissimilarities between neural responses of recognized and non-recognized images for the 13 participants who spontaneously recognized one or more stimuli correctly (eq. 3, cf. Methods).

Figure 3B illustrates the participant-wise dissimilarities of the PRE neural responses to the TPL neural responses. The responses were aggregated into two groups, reflecting whether stimuli were spontaneously recognized before disambiguation or not. We found no evidence for a difference (V = 40, p = 0.6576, g = -0.14 95% CI = [-0.65,0.38]). Bayesian analysis revealed a Bayes Factor of 0.197, indicating evidence against a spontaneous alignment of electrophysiological responses to recognized stimuli before disambiguation towards their photo image counterparts. It is important to note that this evidence is conditional on the limited number of trials available for this analysis.

## Discussion

In the present study, we examined electrophysiological neural responses to two-tone images, and how they align towards the responses to their photo image counterparts after participants have been informed about the stimulus identity via visual disambiguation. We found that electrophysiological responses to two-tone images – in a time window from 0.1 to 0.4 seconds post-stimulus onset – aligned more closely with the response to the photo images after the image content had been learned. This alignment started at around 100 ms post-stimulus onset and peaked between 300 and 400 ms. On the other hand, we found no evidence that the measured neural responses generalized to stimuli of the same, abstract semantic category, as responses to different two-tone stimuli of the same category did not align more to each other after disambiguation. Lastly, we found no evidence for an alignment of neural responses to spontaneously recognized two-tone images and their template counterparts without explicit template exposure.

### Electrophysiological responses to two-tone images converge to their photo image counterparts

Our results show an alignment of electrophysiological responses to two-tone images with those of the corresponding photo images (Fig. 2). This effect is in line with what has been found in fMRI studies focused on the representational alignment of hemodynamic responses ^21^. A positron emission tomography study illustrated such an effect indirectly by demonstrating larger activation of face-selective regions after disambiguation of Mooney face stimuli ^14^. Another recent MEG study demonstrated a better separability of stimulus responses after disambiguation ^24^. However, most recent analyses were likely not able to fully separate the stimulus-specific alignment to similarity alignments which are stimulus-unspecific and attributable to the experimental phase, which we mitigate by subtracting this unspecific component from our data (eq. 1b, cf. Methods, Fig. S8).

While we found positive template alignment values of (re-)learned stimuli, a statistical evaluation of those values for stimuli that were not (re-)learned was precluded by their limited number. By demonstrating this neural alignment in the EEG, the present findings establish two-tone Mooney images as an experimental tool for separating image conceptual content from low-level visual features in neurophysiological paradigms. EEG makes this approach applicable to populations in which fMRI scanning is difficult, such as small children or some patients in a clinical setting. In fact, experiments in young children are usually limited by short experimental times, the absence of a behavioral response, and the interpretability of behavioral proxies such as looking times. While the recognition of Mooney faces (without disambiguation) has been demonstrated in 18 month-olds ^31^ and newborns ^32^ based on looking times, others report recognition only for less degraded or moving two-tone images in younger infants ^33,34^. Thus it is unclear if semantic disambiguation works at this age or in patients with diverse neurological conditions. Thus, experimental paradigms applied to atypical adult cohorts may be less sensitive. For the present study we maximized within-participant sensitivity by presenting 240 repetitions per stimulus and experimental phase, i.e., 2,400 trials per experimental phase and thus 7,200 trials in total, although less can suffice (Fig. S2). Our results demonstrate that two-tone disambiguation can be detected with very brief stimulus presentations: to our knowledge, this is the first demonstration of two-tone disambiguation using a rapid serial visual presentation paradigm with only 250 ms per stimulus. Thus, the observed neural effects can emerge even when each image is available for only a very short period, suggesting that the paradigm may be suitable for populations in which long experimental sessions are difficult to achieve. Moreover, the number of trials can potentially be reduced to discover template alignment (Fig. S2), when less experimentation time is available. An alternative strategy would be to reduce the number of different two-tone stimuli to facilitate successful learning of a small stimulus set instead.

### The neural correlates of template alignment

The strength of evidence for template alignment varied over time in a prolonged time window starting 100 ms after stimulus onset, peaking between 200 and 400 ms, and showing traces even until 800 ms (Fig. 2D). In a previous MEG study, a related effect was reported in a later time window (>500ms) ^24^, although it is not clear if this is driven by a slower stimulation rate (< 0.15 Hz) and longer stimulus presentation (2 s), or based on analytical disparities. Rapid serial visual presentation could, however, artificially shorten this window. For this reason, we deliberately limited our main analyses to the time window of 0.1 to 0.4 seconds.

The strongest contribution to template alignment in and beyond the time window of interest likely stems from lateral occipito-temporal electrodes (Fig. 2E, Fig. S6). Studies using fMRI found template alignment effects in bilateral early visual, lateral occipital and parietal cortices, as well as in bilateral networks distributed across the entire neocortex ^20,21^ including temporal regions known to be relevant for semantic processing ^35,36^. Our electrode-level analysis did not reveal strong template alignment effects on single electrodes pointing towards a more spatially distributed process. It is therefore likely that subtle alignment effects aggregate across a large-scale cortical network, leading to high evidence in the whole brain analysis (Fig. 2A).

### The stimulus, but not the category is encoded after disambiguation

Despite using pairs of stimuli belonging to the same semantic categories (e.g., penguins), our analyses did not reveal that isolated category information is encoded in the neural signal. Although participants did not perform an explicit category perception task, the categories (cow, croissant, hands, human, penguin) were named during disambiguation and understood by the participants. Whether EEG is not able to capture signals related to feature-independent category signals or whether electrophysiological category selectivity requires low-level visual features to be present cannot be answered with the current experiment. Consistent with the latter interpretation it has been shown that EEG decoding performance for image concepts is substantially higher when using visual images than when using their written or spoken counterparts ^37^. Studies using two-tone stimuli in experiments with other techniques such as fMRI or MEG did not investigate category selectivity ^21,24,26^.

The present experiment was designed to perfectly equalize the low-level visual features to analyse template alignment, by applying the same physical stimuli and only varying the perceptual state (recognized vs. not recognized). However, low-level visual features naturally still differed within and between categories, so two two-tone cows could share more low-level visual features than a two-tone cow and a two-tone penguin. Likewise, the similarity of low-level visual features inherently differs between two-tone and photo images, as photo images contain richer low-level visual features. As a result, only the direct statistical comparison of category selectivity between PRE and POST phases is truly unbiased by low-level features, as they also mitigate potential biased offsets of category selectivity values of each phase. In a pixel-wise comparison, however, distances between and within categories were evenly distributed across photo and two-tone images (Fig. S7).

For the analysis of category selectivity, the within-category term in equation 2 is likewise bound by categories which were both (re-)learned by each participant, and thus does not fully capitalize on the large number of presented stimuli, limiting the power of the analysis. On a similar note, we were bound by a tradeoff between the number of total stimuli, stimuli per category and the number of categories. The number of total stimuli is inherently limited, and we found 10 reasonable in this experimental context for healthy adults which limited our exemplars to two per category. More stimuli per category bear the risk of more cross-talk during the PRE phase (i.e., recognizing one penguin can support recognizing another), and limiting the amount of categories similarly limits the generalizability. Furthermore our images were not selected for ‘cleanness’ of exemplar presentation further limiting the interpretability. Future studies tailored to unveil category selectivity should aim for a more diverse stimulus set and, e.g., an interleaved presentation of stimuli from different experimental phases (PRE, TPL & POST) ^24^.

### Low-level image features dominate the electrophysiological signal

As shown in Figures 2C, S4, & S5, electrophysiological responses were more similar, when the physical stimulus was the same (PRE vs. POST), compared to when the semantic concept was the same (POST vs. TPL). This difference illustrates the large influence of low-level image features on the electrophysiological signal compared to the more subtle influence of the semantic concept, at least in the observed time window spanning 0.1 – 0.4 seconds. This finding is particularly important when interpreting the results reported in studies in which category-level decoding was performed on electrophysiological responses elicited by photographic stimuli ^5,25,38,39^. Only a few studies have analytically tried to quantify or control for low-level features ^9,40–45^. Above-chance category decoding reported in such studies may therefore be partly driven by low-level image features rather than by semantic category information alone. A rapid serial visual presentation paradigm, as used in the current study, could lead to an artificial shift of this balance towards low-level feature processing, as downstream semantic processing steps might be partly attenuated by the dense visual input, although it is unlikely that subsequent stimuli fully curtail semantic representations, as they do also not curtail the general neuronal representation of images ^3,27^.

If an emergent semantic signal existed for particular categories, it might require less degraded sensory input. Other techniques, such as fMRI or MEG, might be less dominated by the low-level visual signal. For instance, magnetoencephalographic responses to POST-disambiguation stimuli are much closer to those of their photo image counterparts than to their PRE-disambiguation versions ^24^, contrary to what we show using EEG. Furthermore, fMRI studies show template alignment even within the early visual cortex, where low-level features should dominate the neural responses ^21^.

Currently, there is no exhaustive account of which image properties can be considered as low-level, nor whether these properties can be fully parameterized independently of semantic category. Rather than forming discrete representational levels, low-level visual statistics and semantic information may instead lie on a continuum of increasingly abstract image regularities, complicating attempts to isolate purely semantic neural representations ^44,46–50^.

### Neural alignment follows template exposure, not concept understanding

Finally, we did not find spontaneous alignment between neural responses to recognized stimuli of the PRE-disambiguation phase with their photo image counterparts of the TPL-phase. However, the power for detecting spontaneous alignment is more limited, as only ∼12% of all PRE and TPL trials across participants could be recruited for the analysis, compared to ∼85% of trials that entered the template alignment analyses (Table S1, Fig. 2B). Furthermore, we did not systematically ask participants, from which timepoint onwards they recognized the image in the PRE phase. Stimuli recognized early could lead to stronger effects than stimuli recognized only in the third PRE block. Lastly, whether a particular two-tone image is perceived in a more holistic or more feature-dependent manner is prone to individual differences between participants ^51,52^ and cannot be inferred from the acquired participant responses. Future experiments might include additional behavioral tests to further subdivide stimuli based on participant- and stimulus-specific cognitive strategies, to quantify participant engagement and attention, and to better individualize diverse cognitive strategies across the experiment. On the same note, introducing a non-disambiguated control condition could strengthen the link between representational alignment and recognition while at the same time controlling for time-varying cognitive states.

The absence of alignment in spontaneously recognized images, however, also raises the possibility that the observed effect is at least partly specific to exposure to the visual template, rather than reflecting semantic knowledge per se. Repeated exposure to the template may facilitate subsequent processing via perceptual learning and increased familiarity, i.e. once the interpretation of an ambiguous image has been established, subsequent processing may become more efficient because the sensory input can be matched against an existing representation. One way to disentangle these possibilities in future work would be to provide prior knowledge about the image content without exposing participants to the corresponding visual template – for example, by verbally informing participants of the object’s identity ^16^. If semantic information alone were sufficient to produce the subsequent alignment, this would support an interpretation in terms of semantic enrichment; conversely, a stronger dependence on template exposure would suggest that perceptual reinstatement, template-specific learning, or the coupling between semantic and perceptual representations contributes to the effect. We did not include such a verbal-cue condition in the present experiment, and therefore cannot distinguish these possibilities here.

The absence of spontaneous alignment is also consistent with the absence of a strong category signal. An emergent category signal could rather facilitate spontaneous alignment, because the category is more easily perceived spontaneously than the stimulus-specific peculiarities such as edges and cluttering not related to the central object. One possible interpretation is that recognition of two-tone images does not necessarily entail reinstatement of a fully specified, stimulus-faithful representation, but instead reflects a partially specified perceptual state in which different levels of representational detail can be available to varying degrees. According to this view, recognition may rely on sufficient information for object-level identification without requiring recovery of the full set of image-specific features that distinguish one stimulus from another. This interpretation is consistent with literature on image disambiguation, of which two-tone perception is a specific case, suggesting that perceptual outcomes vary depending on available priors and learned templates, with systematic differences regarding the degree to which fine-grained visual detail is recovered ^53,54^. More broadly, work on interactive feature processing in perceptual organization suggests that holistic or global interpretations can be formed even when local feature analysis remains incomplete or unstable ^55^.

Several additional possibilities are compatible with the present findings. First, spontaneous recognition may involve semantic access that is relatively weak, transient, or perceptually underspecified. For example, Ishikawa and Mogi ^15^ reported reversals between correct and incorrect identifications of two-tone images, suggesting that the perceptual interpretation of such stimuli can remain unstable even when an interpretation has been reached. Furthermore, even subliminal presentation of templates has been shown to facilitate Mooney disambiguation ^56^ demonstrating that top-down information can influence perceptual processing even when the information itself is not consciously accessible.

Second, template exposure may strengthen or stabilize an interpretation that was already partially available, making it sufficiently robust and temporally consistent to produce detectable neural alignment. Template exposure could also reduce perceptual uncertainty or selectively reweight diagnostic visual features via attention allocation, thereby producing a more stable perceptual organization of the image. Alternatively, the template may establish a stimulus-specific association between the particular visual configuration and its interpretation rather than a conceptual one in line with the findings reported by Hachisuka et al. ^57^, such that subsequent processing benefits from memory-based or top-down facilitation. These possibilities are not mutually exclusive, and the present data do not allow us to distinguish between them. Importantly, a dissociation between semantic access and reinstatement of detailed perceptual information has also been demonstrated specifically with two-tone images. Hegdé and Kersten ^58^, using fMRI, reported partially distinct neural signals associated with memory for the original photograph and with disambiguation of the corresponding two-tone image, with neither signal reliably predicting the other. This provides converging evidence that successful disambiguation of a two-tone image need not entail reinstatement of the full visual representation associated with the corresponding photograph. However, the extent to which this finding can be directly generalized to the present experiment is limited. Hegdé and Kersten used only two object categories and a different procedure for assessing disambiguation, making their design particularly suited to examining the contribution of visual memory to a more constrained recognition task.

Together, these findings support the idea that semantic access and detailed perceptual reinstatement are at least partially dissociable, rather than necessarily co-occurring. At the same time, the present results cannot determine whether template exposure introduces semantic information that was not previously available, strengthens or stabilizes a weakly available interpretation, enhances the perceptual features supporting that interpretation, or establishes a stimulus-specific memory association.

## Conclusion

In summary, our results suggest that EEG signals elicited by two-tone images carry information about learned content, as demonstrated by a stimulus-specific alignment between neural responses to disambiguated two-tone images and their corresponding photo templates. This template alignment emerges rapidly after stimulus onset and persists over time, indicating that learned information about image content can systematically modulate electrophysiological responses even in the presence of highly degraded visual input. At the same time, our findings provide neither evidence for a category-level representation of semantic information in the EEG signal nor for spontaneous alignment between neural responses and their corresponding templates prior to explicit visual disambiguation. Instead, the data suggest that neural similarity structure is dominated by low-level visual properties, with conceptual effects representing a more subtle and partially separable cognitive contribution within the same signal space.

## Methods

### Participants

17 healthy adults (13 female) with a median age of 24 years (range: 20 – 34 years) were invited to participate in an EEG session. All participants had normal or corrected-to-normal vision and gave written informed consent. The study was approved by the Ethics Committee of the Medical Faculty of the University of Leipzig, Germany (402/23-ek). Data acquisition took place at the Max Planck Institute for Human Cognitive and Brain Sciences in Leipzig, Germany.

### Stimulus creation

Two-tone images were generated from natural photographs for five categories: cows, croissants, hands, humans, and penguins. Most images (cows, croissants, penguins) were derived from the THINGS database ^59^. Human bodies and hands are highly salient stimuli, but no sufficiently ambiguous, i.e. not spontaneously recognisable two-tone images could be generated from the images in the database as their characteristic contours and structures kept these highly salient stimuli easily disambiguable. Accordingly, copyright-free images from the internet were used instead for hands and human bodies. See Table S2 for more information on the original source of all used images. Input images were first converted to grayscale. Images were resized to a fixed resolution of 400 × 400 pixels. Grayscale images were then spatially smoothed using a Gaussian filter, with the standard deviation of the Gaussian kernel (σ) between 2pxs and 7pxs controlling the degree of smoothing. Smoothing kernels were manually selected on a per-image basis to achieve an optimal trade-off between image distortion and the ability to associate the two-tone image with its original template image after disambiguation. Smoothed images were binarised using global intensity thresholding. For each image, a single threshold value was computed using Otsu’s method ^60,61^, and pixel intensities were mapped to black or white accordingly. The resulting binary images were scaled to an 8-bit intensity range (0–255) and saved as grayscale JPEG files.

We created ten different two-tone images (Fig. S1), comprising five semantic categories with two distinct images per category (cow, croissant, hands, human, penguin). From an initial screening of approximately 500 images, ∼50 additional candidate images and categories were evaluated in pilot experiments. The final set of stimuli was selected based on a low spontaneous recognition rate prior to template exposure and the subjective ease with which the two-tone images could be associated with their corresponding template images.

### Experimental design

The experiment included three main phases: PRE-disambiguation (PRE), Template (TPL), and POST-disambiguation (POST). Each phase comprised 2,400 stimuli distributed across three blocks (Fig. 1), leading to a total of 7,200 stimulus presentations to maximize power within participants.

During the PRE-disambiguation phase, two-tone images were presented at a rate of 2.5 Hz (250 ms stimulus duration, 150 ms inter-stimulus interval). Participants were instructed to attend to the screen and focus on the stimuli without performing an explicit task. Each block consisted of 80 repetitions of each stimulus presented in random order allowing for repetitions of the same stimulus, resulting in a total of 800 stimulus presentations per block and a block duration of 5 min 20 s.

After completing the final PRE block, participants were briefly (250 ms, with option for repetition) shown each single two-tone image and were asked whether they had recognized the content depicted in the image in the previous blocks and, if so, what they believed it depicted. Participants were unaware of any stimulus category labels, and freely described either the recognized scene or object they saw or think they saw. Based on the participants’ responses, the experimenter classified each stimulus as not recognized, falsely recognized (i.e., a reported concept that was clearly distinct from the true underlying image), or spontaneously recognized (i.e., correct identification of the image content, Table 1).

Next, each stimulus was disambiguated using a fade-in/fade-out procedure. Specifically, the two-tone image was first presented, after which its opacity was gradually reduced to reveal the underlying grayscale photo template image, and subsequently increased again to restore the two-tone image. While the template image was fully visible, a computer-generated voice verbally provided the category label (cow, croissant, hands, human, penguin, in German) to facilitate recognition and to ensure that participants understood the intended image content. One automated disambiguation cycle lasted 5 seconds (1 s two-tone, 1 s fade-in, 1 s template, 1 s fade-out, 1 s two-tone), and was performed once for each stimulus within each disambiguation block. Disambiguation blocks were presented once prior to the template phase and again before each POST block. Following each automatic disambiguation block, participants were asked whether they experienced difficulties recognizing any of the stimuli. If so, an additional manual disambiguation cycle was conducted, during which participants could control the speed of the transition between two-tone and template images.

Subsequently, the grayscale photo template images were presented in three TPL blocks. Stimulus timing and presentation parameters were identical to those used in the PRE phase. Participants were again instructed to attend to the stimuli without performing an explicit task.

The subsequent POST phase comprised three blocks. These blocks were identical to the PRE blocks, except that each POST block was preceded by a disambiguation block to refresh participants’ memory of the ten two-tone images and their underlying template.

After completion of the final POST block, participants were again briefly presented with each two-tone image and asked to report its identity. Based on these responses, the experimenter classified each stimulus as learned (not recognized during PRE, but correctly identified after disambiguation), re-learned (falsely recognized during PRE, but correctly identified after disambiguation), or not (re-)learned (not or falsely recognized during PRE and still not correctly recognized after disambiguation, Table 1).

The entire experiment lasted approximately 90 minutes including breaks, and required an additional ∼30 minutes for EEG cap preparation. Participants were allowed to take breaks ad libitum between blocks. Stimuli were presented at 5.3° of visual angle on a uniform grey background. This relatively small stimulus size was chosen based on pilot experiments indicating that this setup facilitated perception of the images. Images in each experimental block were displayed in random order.

### Data acquisition

Participants were seated in front of a computer screen in a dimly lit room. EEG was recorded using active electrodes (Brain Products) arranged in a custom montage comprising 34 electrodes: 31 scalp EEG electrodes (Fz, F3, F7, F9, FC5, POz, C3, T7, TP9, CP5, P7, P3, Pz, O1, O2, P8, P4, CP6, TP10, T8, C4, Oz, FC6, F10, F8, F4, Fp2, FC4, CP4, FC3, CP3), one electrooculogram (EOG) electrode situated on the right cheek, an online-reference electrode at FCz, and a ground electrode at Fp1. No online filter was applied.

EEG signals were recorded using BrainVision Recorder (version 1.22.0001, Brain Products) at a sampling rate of 1000 Hz. EEG caps (Easycap) were fitted to the participants’ heads, and a custom-made conductive gel (hydroxyethyl cellulose, propylene glycol, sodium chloride, and water) was applied until electrode impedances were reduced to approximately below 20 kΩ. Electrodes exhibiting poor signal quality (e.g., flat signal, persistently high impedance, or excessive noise) were documented during recording and excluded from further analysis prior to preprocessing. The experimenter continuously surveilled participant engagement via a video camera providing a frontal, close-up view of the participants face.

### Data preprocessing

Data processing was performed using Bash, Python (v. 3.11.14), and R (v. 4.3) on a high-performance computing cluster at the Max Planck Computing and Data Facility (Garching, Germany). Electrophysiological data were preprocessed using MNE (v. 1.11.0; ^62^).

First, channels exhibiting excessive noise were manually identified and removed. The raw data were then resampled to 250 Hz, anti-aliasing was applied as an intrinsic part of the resampling procedure, with the signal band-limited to the new Nyquist frequency of 125 Hz before the sampling rate was reduced. In participants for whom the necessary electrodes were available, two artificial EOG channels were constructed: (1) by subtracting the voltage of the physical EOG electrode located below the right eye from electrode Fp2 (above the right eye), and (2) by subtracting the signals of electrodes F9 and F10 ^39,63^. The physical EOG channel was subsequently removed from the dataset.

The continuous time series was high-pass filtered at 0.1 Hz and low-pass filtered at 30 Hz using a linear finite impulse response, zero-phase non-causal band-pass filters with a windowed time domain design method (firwin), and a Hamming window with 0.0194 passband ripple and 53 dB stopband attenuation, with transition bandwidths of 0.1 and 7.5 Hz and -6 dB cutoff frequencies of 0.05 and 33.75 Hz and a filter kernel duration of 33 seconds. To further identify and correct noisy channels and to compute a reference, the pyprep pipeline ^64^ including the RANSAC algorithm ^65^ was applied. Identified bad channels were interpolated, and all channels were re-referenced to the resulting robust average reference. Independent component analysis (ICA) was performed using the Picard algorithm ^66^ on a copy of the data that had been high-pass filtered at 1 Hz ^67^. Twenty independent components were estimated with a maximum of 500 iterations. Each component was correlated with the available artificial EOG channels. Components were classified as ocular artifacts if the correlation exceeded a threshold determined via adaptive z-scoring. The resulting ICA unmixing weights were then applied to the bandpass-filtered data, by reconstructing the component time courses, zeroing the components deemed as ocular artifact, and recomposing the sensor-space data from the remaining components.

Data were subsequently epoched using a pre-stimulus baseline of 400 ms (corresponding to one stimulus cycle, e.g. ^68^) and a post-stimulus onset interval extending to 800 ms after stimulus onset (corresponding to two stimulus cycles). Linear detrending was applied across the entire epoch, followed by mean-centering using the pre-stimulus baseline interval. Additional artifact correction was performed at the epoch level using the autoreject package ^69,70^ v. 0.4.3. Hyperparameters were configured to reject, but not interpolate, contaminated epochs, as channel interpolation had already been performed at an earlier preprocessing stage. Rejection thresholds were estimated using 5-fold cross-validation.

Artifact rejection resulted in slight imbalances in the number of trials, potentially biasing distance estimates. We therefore equalized trial counts across conditions within participants via undersampling – i.e., by dropping the chronologically latest surplus epochs.

### Dissimilarity estimation and aggregation

We estimated Mahalanobis distances as dissimilarity measures between epochs belonging to the different conditions using all electrodes and a post-stimulus onset time window from 0.1 to 0.4 s. Mahalanobis distance was selected because it takes the covariance between the spatiotemporally dependent features into account. The window was chosen to cover nearly one stimulus cycle, starting shortly after stimulus onset when a neural response can be expected, and ending at the next stimulus onset. The resulting channel × time point data matrices were transformed into a single feature vector ^71,72^, and pairwise neural dissimilarity between conditions was quantified. All possible pairwise distances were computed, and the respective distance values were extracted for evaluating our hypotheses (see below). We used a shrinkage estimator to compute the (inverse) covariance matrix ^73^ across all trials of a participant. Mahalanobis distance estimation was conducted separately for each participant to account for the different noise levels of each electrode and each participant. To further account for different noise levels between participants, all inference is done on either difference values – calculated within-participant – or using paired statistical tests.

#### Primary hypothesis: Template alignment (TA)

To test whether neural responses to two-tone images became more similar (less dissimilar) to responses to their corresponding template images after disambiguation, we computed, for each participant (p), the average difference in dissimilarity between PRE vs. TPL and POST vs. TPL conditions (paired for each stimulus) across all learned and relearned stimuli (L). Template alignment was quantified as:

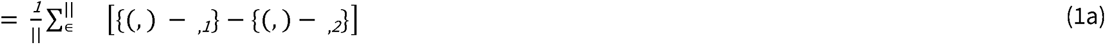

To exclude stimulus-unspecific effects, we subtracted a participant-and phase-comparison-dependent constant,. Whereas the effect of interest was calculated by pairing the same stimuli, it cannot be avoided catching also unspecific effects that can be attributed to the experimental phase. For example, if neural responses to all PRE stimuli share (or lack) a common signal, the dissimilarity metric between any unpaired PRE and TPL stimulus (e.g., cow_02_PRE vs. penguin_06_TPL) will be higher, than the dissimilarity metric between any unpaired POST and TPL stimulus (e.g., cow_03_POST vs. croissant_03_TPL). The template alignment result will therefore be biased by unspecific effects that just differentiate experimental phases. As a solution, we systematically calculated all possible distances (i.e., the Cartesian product) of all possible (paired and unpaired) comparisons, while omitting paired comparisons, and averaged across those:

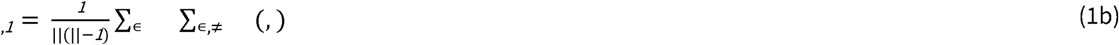

_,2_ is then calculated analogously for the dissimilarity between POST and TPL. A resembles the set of all 10 stimuli. We evaluated the effects of eliminating these constants from the analyses and its bias of the template alignment metric in Figure S8.

To further evaluate temporal properties of template alignment, we repeated the same analysis for small consecutive time windows (12 milliseconds, corresponding to 3 samples of the preprocessed epochs), using the data of all electrodes. We also explored the spatial topography of template alignment by applying the same formula to data from the previous time window of interest (0.1 s – 0.4 s) but for each electrode separately. Lastly, we repeated the electrode-wise analysis for consecutive time-windows of 100 ms.

#### Secondary hypothesis 1: Category selectivity (CS)

To assess whether image category information, beyond individual stimulus identity, was encoded in neural responses after learning, we computed, separately for the PRE, POST, and TPL phases, neural dissimilarities between pairs of images of the same phase belonging to the same category (within-category pairs, W) or to different categories (between-category pairs, B). Only image pairs for which both stimuli were classified as learned or re-learned were included.

Category selectivity was defined as the difference between between-category and within-category distances, computed for each participant (p) as:

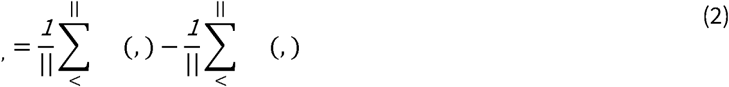

Category selectivity was calculated similarly for the other phases (POST and TPL), and the difference in category selectivity between phases is computed using paired tests.

#### Secondary hypothesis 2: Spontaneous alignment (SA)

To examine whether spontaneously recognized two-tone images elicited neural responses more similarly to their corresponding template images prior to disambiguation, we compared PRE vs. TPL dissimilarities between spontaneously recognized stimuli (S) and stimuli that were not recognized (N). For each participant (p), spontaneous alignment was defined as:

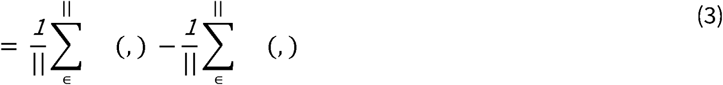

Multidimensional scaling on distance values. To visually illustrate how the stimulus-wise distance values between the three experimental phases are related, we computed a metric multidimensional scaling model with only one dimension. Therefore, we used the mean of all learned and re-learned stimuli’s raw Mahalanobis distance values for each combination of experimental phases (PRE vs. POST, PRE vs. TPL, POST vs. TPL), averaged across participants, and entered them to a metric multidimensional scaling function (R, stats::cmdscale v. 4.3.3) with default hyperparameters. The first dimension explained 66.5% of variance in the data. We further repeated the same process for each individual participant.

### Statistical analyses

To test whether template alignment (TA), category selectivity (CS), or spontaneous alignment (SA) were greater than zero, we computed one-tailed Bayes Factors (BFs) using the R BayesFactor package (v. 0.9.12-4.7). Cauchy priors with a scale parameter of 0.707 were used which provide rather conservative prior distributions on the effect sizes. Reported BFs > 1 quantify the evidence in favor of the hypothesis that the respective effect is greater than zero. BFs < 1 quantify the evidence for an effect smaller or equal to zero. The classification of Kass & Raftery ^74^ was used to verbally qualify evidence based on BFs, with a BF > 3 considered evidence, and a BF > 20 considered strong evidence (analogous for the inverses). For timepoint-wise and electrode-wise analyses, the same criteria were applied ^29^. We either tested if 0, 0, or if 0. In complementary frequentist analyses, we computed one-tailed Wilcoxon signed-rank tests, using an arbitrary critical alpha level of 0.05. For multiple paired variables we applied Friedman tests. Effect sizes and corresponding confidence intervals were calculated using Hedges’ g, to account for a relatively small sample size.

### Perceptual similarity

To quantify perceptual image similarity, we computed (atanh-transformed) Pearson correlations between the pixel intensities of all pairs of images ^75^, separately for photos and two-tone images (10 images each; 45 pairs per image type, of which 5 belonged to the same category). Because each image contributes to several pairs, the pair-wise similarities are not independent, which rules out conventional parametric tests. We therefore used exact permutation tests that permute labels at the level of images, preserving the dependency structure of the similarity matrix. To test whether images of the same category were more similar than images of different categories, we compared the mean within-category and between-category similarity under all 945 possible assignments of the 10 images to five categories of two exemplars. The interaction with image type was tested in the same way on the per-pair difference between two-tone and photo similarities. To compare image types, we exchanged the photo and two-tone version of each image under all combinations, using cross-version correlations for pairs whose images were assigned to different versions. All tests were two-sided; p-values were Holm-corrected within families of tests.

## Data availability

The raw data collected for this study is available through a public link on Zenodo (https://doi.org/10.5281/zenodo.20037230).

## Code availability

All code used for data analysis is available from GitHub (https://github.com/SkeideLab/MOONEY_adults). The experimental paradigm is available for download from Pavlovia (https://gitlab.pavlovia.org/kesslerr/MOONEY_adults), and code to generate Mooney images is available from GitHub (https://github.com/Algebreaker/MooneyGeneration).

## Funding statement

This work was supported by the German Research Foundation (DFG Heisenberg Program Grant 433758790 awarded to M.A.S.) and the Jacobs Foundation (Research Fellowship awarded to M.A.S.). The funders had no role in study design, data collection and analysis, decision to publish or preparation of the manuscript.

## Acknowledgements

Not applicable.

## Author contributions

Conceptualization: RK, JF, MAS; Formal analysis: RK; Funding acquisition: MAS; Investigation: RK, JF; Methodology: RK, JF; Project administration: RK, JF, MAS; Ressources: RK, MAS; Software: RK; Visualization: RK; Writing: original manuscript: RK, JF; Writing: review & editing: RK, JF, MAS

## Ethics declarations

The authors declare no competing interests.

## Supplementary Information

**Fig. S1.**
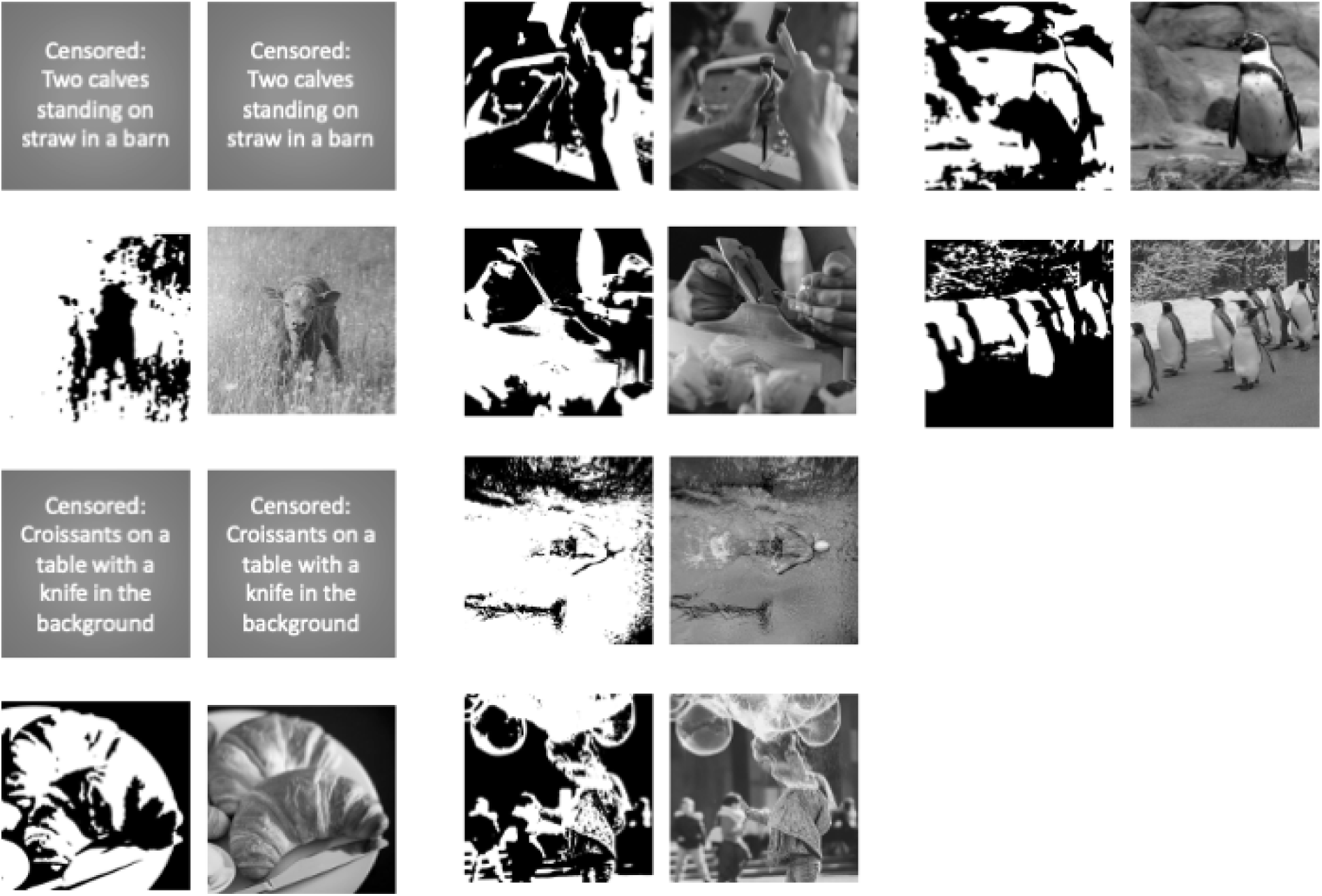
Stimuli. Ten images of five categories (cow, croissant, hands, human, penguin) were converted to greyscale (right columns), smoothed, and binarized to create two-tone images (left columns). See Table S2 for image sources. Images with unclear copyright information have been censored.

### Sequential trial accumulation analysis

For each participant, different percent of artifact-free trials were used to evaluate template alignment on the group level based on the learned and re-learned images. Therefore, we first equalized trial counts per condition within participants based on the condition with the minimum number of artifact-free trials (Fig. S2C). We then used several fractions of the available trials to repeat our main analysis regarding template alignment. The chronologically first trials were used to reflect their order in the experimental design. Results illustrate that with about 25% of trials, a Bayes Factor of 3 wa exceeded, and with about 70% of trials a Bayes Factor of 10 was exceeded (Fig. S2B). However, the effect sizes still increased with higher trial counts (Fig. S2A).

**Fig. S2.**
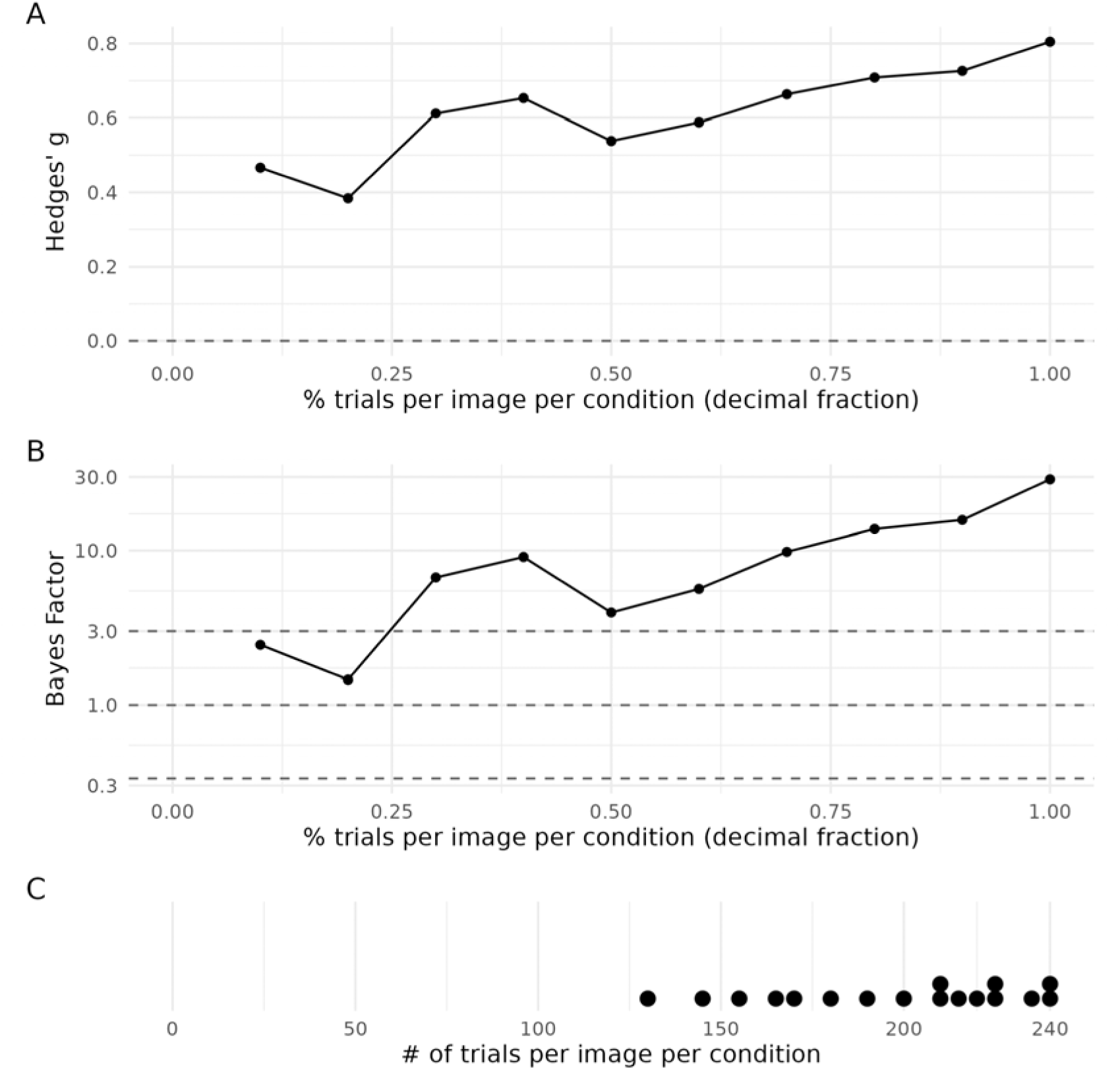
Sequential trial accumulation analysis on template alignment. For each participant, different fractions of artifact-free trials were used to evaluate template alignment on the group level. Hedges’ g (A) and Bayes Factors (B) were calculated as a function of percentage artifact-free trials (x-axes). The dotplot (C) illustrates the number of artifact free trials (minimum across conditions) per participant. This minimum was used and surplus trials of other conditions were discarded. The percentage of artifact free trials was then related to this number of trials.

### Design sensitivity analysis

A Bayesian sensitivity analysis was conducted to characterise the effect sizes that the present design was able to detect. We used the BFDA package ^1^ with N = 17, a one-sided default Cauchy prior on effect size (r = 0.707, truncated at zero), and Bayes factor thresholds of 3 and 10. The design would yield BF > 3 and > 10 with a probability of 80% for a true effect greater than 0.71 and 0.89, respectively (Fig. S3). The design was therefore sensitive to large effects, whereas small to moderate effects would frequently have remained inconclusive. Note that this analysis is independent of the observed data and reflects a property of the design rather than a restatement of the obtained Bayes factor.

**Fig. S3.**
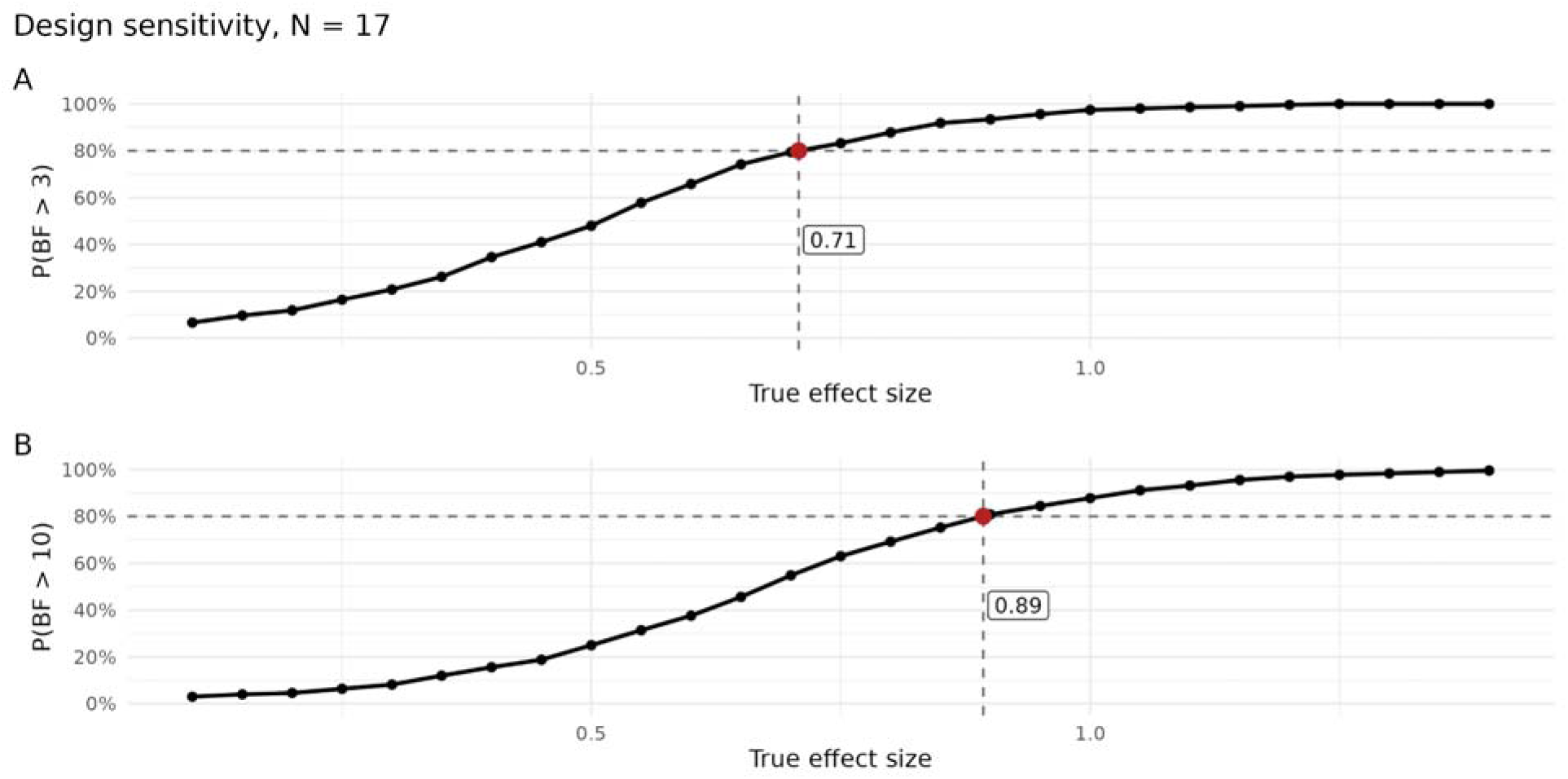
Design sensitivity for template alignment. The probability of hitting a critical Bayes Factor boundary of (A) or 10 (B) is illustrated on the y axes for different true effect sizes (x axes). Dashed horizontal lines indicate the probability of exceeding the critical Bayes Factor with a probability of 80%. Dashed vertical lines and associated red dots indicate the intercept for each Bayes Factor boundary.

**Fig. S4.**
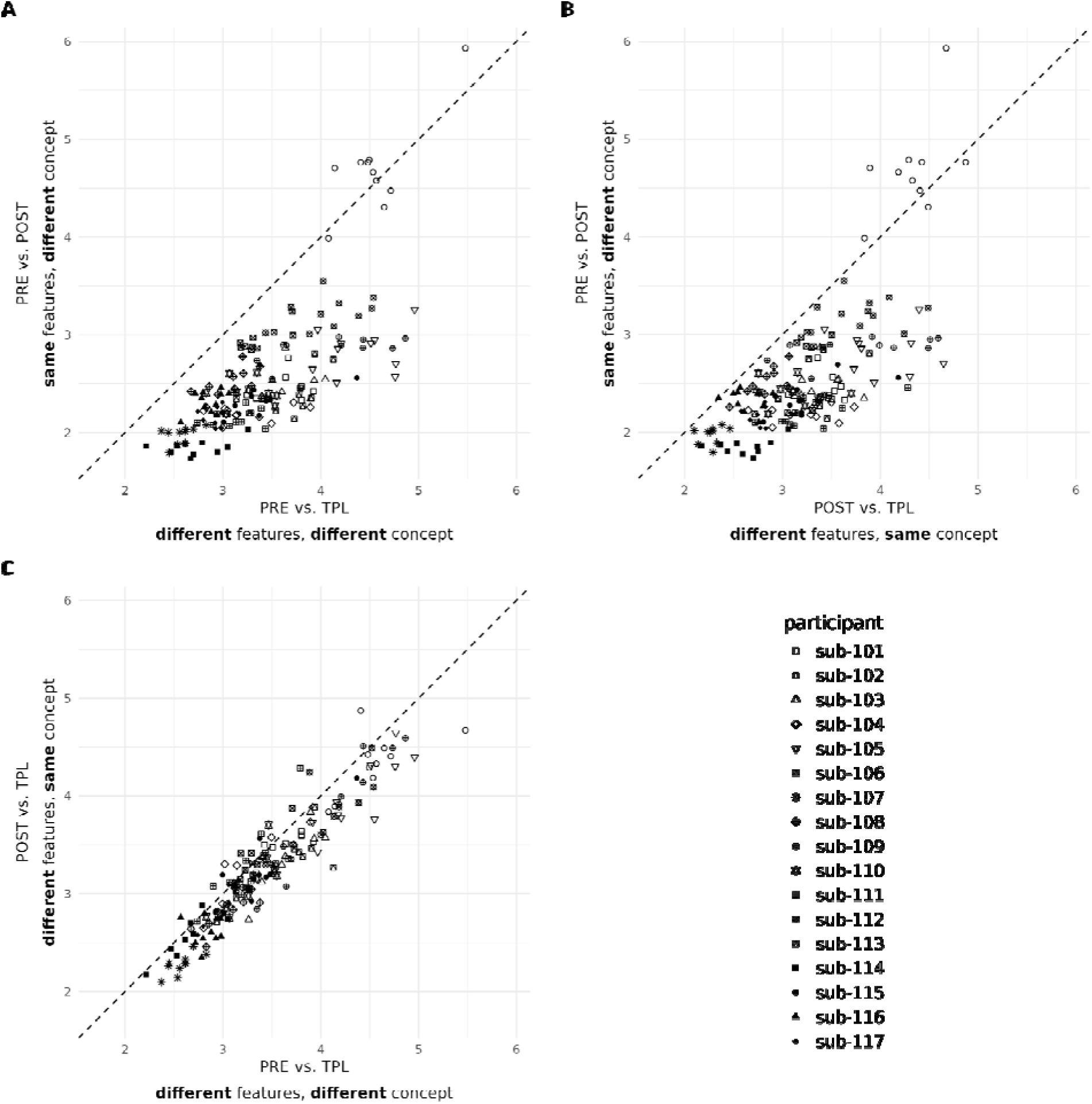
Comparison of Mahalanobis distances between experimental phases. The stimulus-and participant-wise Mahalanobis distances are plotted for different comparisons of experimental phases. Subjects ar indicated by different shapes. The dashed diagonal is the identity line. A: PRE vs. POST against PRE vs. TP distances. The comparisons differ only in the low-level features. B: PRE vs. POST against POST vs. TPL distances. The comparisons differ in low-level features and semantic concept. C: POST vs. TPL against PRE vs. TP distances. The comparisons differ only in the semantic concept. The distribution shows smaller distances in POST vs. TPL, indicating template alignment. The graph offers a different view to the same data from Figure 2. All stimuli are shown, irrespective of their group, i.e., (re-)learned, spontaneous, and not (re-)learned. Th electrophysiological response is more similar (less dissimilar) between PRE and POST conditions than either of them with the TPL condition in most participants and stimuli, indicating that low-level image features are a stronger driver for dissimilarity than semantic concept. The filtered data points for learned and re-learned stimuli per participant formed the basis for the multidimensional scaling approach of Figure 2C & Figure S5.

**Fig. S5.**
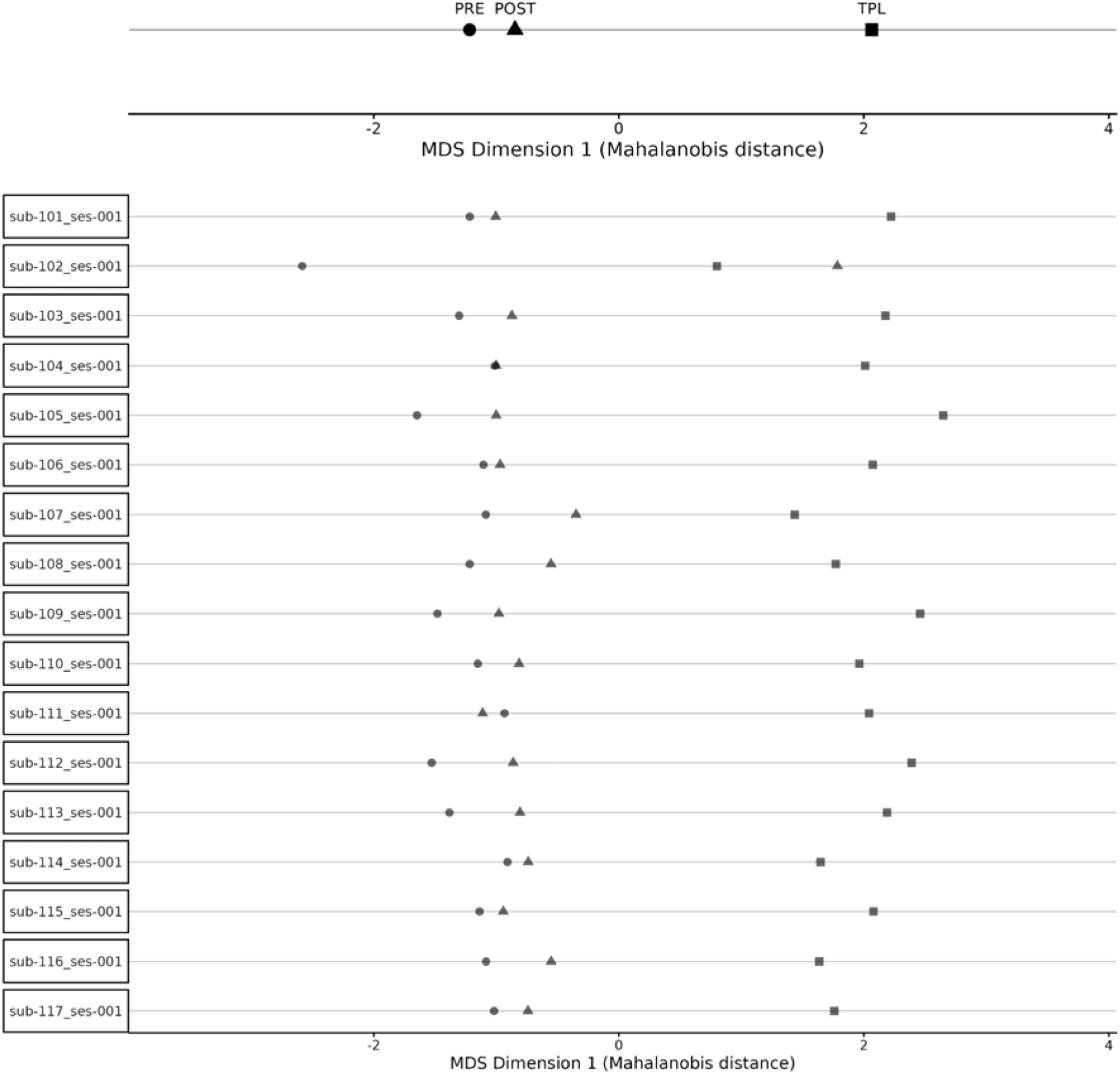
Multidimensional scaling. Metric multidimensional scaling was performed using the pairwise distances between the three experimental phases (PRE, TPL, POST) for the group (upper panel) each individual participant (lower panel). Note that because the direction on the axis is arbitrary, the sign was flipped for each participant so that the TPL condition is always located right of the PRE condition. 15 out of 17 participants expressed the ordering PRE – POST – TPL.

**Fig. S6.**
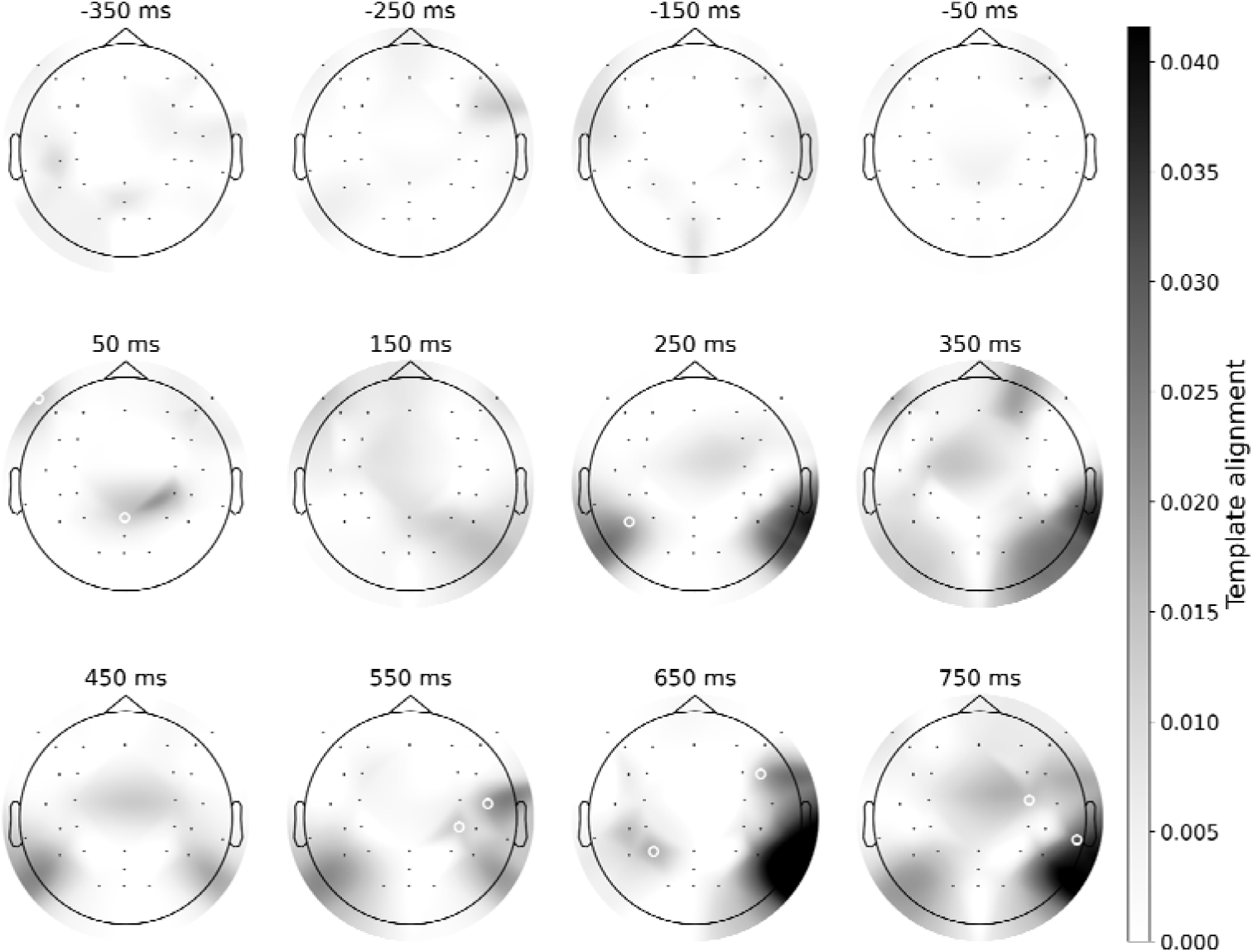
Spatio-temporal template alignment. For each 100ms time window (centered at the time indicated in each facet title), and for each electrode individually, the mean template alignment across participants was computed (luminance-coded). White circles indicate electrode-time-window combinations exceeding a Bayes Factor of 3. Note the inflated rate of false positive evidence due to the large number of tests.

**Fig. S7.**
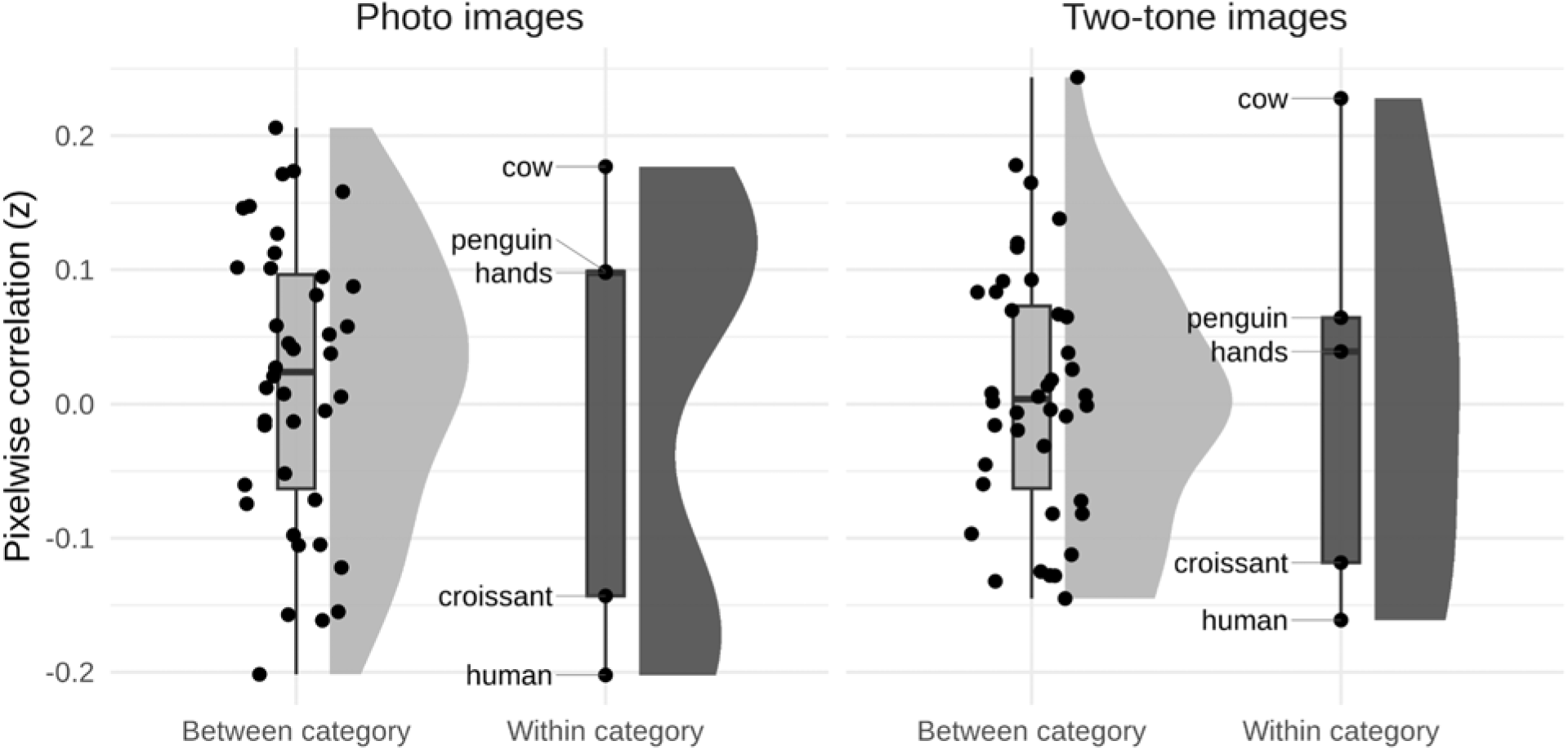
Perceptual similarity within and across stimulus categories. Between each pair of images, the perceptual similarities as the (atanh-transformed) correlation coefficients (y axis) were computed, either within category (black) or between categories (grey). Within category similarity values are labeled with the category. Boxplot corpus illustrates 0.25, 0.5, and 0.75 quantiles. Whiskers span all data points within 1.5 times the inter-quantile-range. We found no evidence for higher similarities in photo (left panel) vs. two-tone (right panel) images (all p > 0.05, two-sided permutation tests). Furthermore, perceptual similarities were not different within or between categories (all p > 0.05, two-sided permutation tests).

### Removing stimulus-unspecific effects

To compute template alignment values, we introduced participant-specific constants, (eq. 1b) in order to subtract those from the distance values of each experimental phase with the template phase within equation 1a (cf. Methods). Without those constants, the (difference of) distance measures of the neurophysiological responses would be biased by stimulus-unspecific effects. Figure S8 illustrates the Bayes Factors of the time-resolved template alignment values when omitting those constants.

Each epoch includes 3 stimulus-cycles. Stimuli were presented at -400 ms, 0 ms, and 400 ms. For any analysis, epochs were ordered based on the label of the stimulus presented at 0 ms, whereas the choice of the other two stimuli, one preceding and one succeeding, were random. More specifically, when the trials were sorted for the analyses and the stimulus at 0 ms always showed cow_02 (PRE, POST, or TPL), the stimuli at -400 ms and 400 ms comprise random (other or same) stimuli, as the presentation was randomized. When not including, in equation 1a, those template alignment values become clearly positive, even with unordered stimuli (first and third peak), indicating similarity differences that are specific to the experimental phase rather than to this exact stimulus and the experimental phase. More specifically: Figure S8 shows three peaks: Peak #1 (from left to right) is only sensitive to the experimental phase, because stimuli are unordered. Peak #2 is sensitive to the experimental phase and the stimulus, as stimuli in this time window are ordered. Peak #3 can theoretically be both, either an effect of the experimental phase, or also an effect of the preceding stimulus which is again ordered. When introducing the constant to the equation, the unspecific peak #1 completely vanished from the baseline period (compare Figures 2D and S8). Peak #2 lost its stimulus-unspecific parts, reduced in amplitude and elongated. Peak #3 as a separate entity also vanished.

The reason for the stimulus-unspecific peaks can be manifold. On the one hand, if the PRE trials were more often accompanied by face pareidolia, face specific components could be encoded in the neuronal signal ^2^. Therefore, the PRE trials express some “extra” component, and the first term in equation 1 gets larger if neglecting the constant, leading to spurious positive values for template alignment. On the other hand, if the POST and TPL trials were accompanied by a “recognition” signal – as in both phases stimuli were recognized – it renders again the first term in equation 1 large, leading to spurious template alignment.

**Fig. S8.**
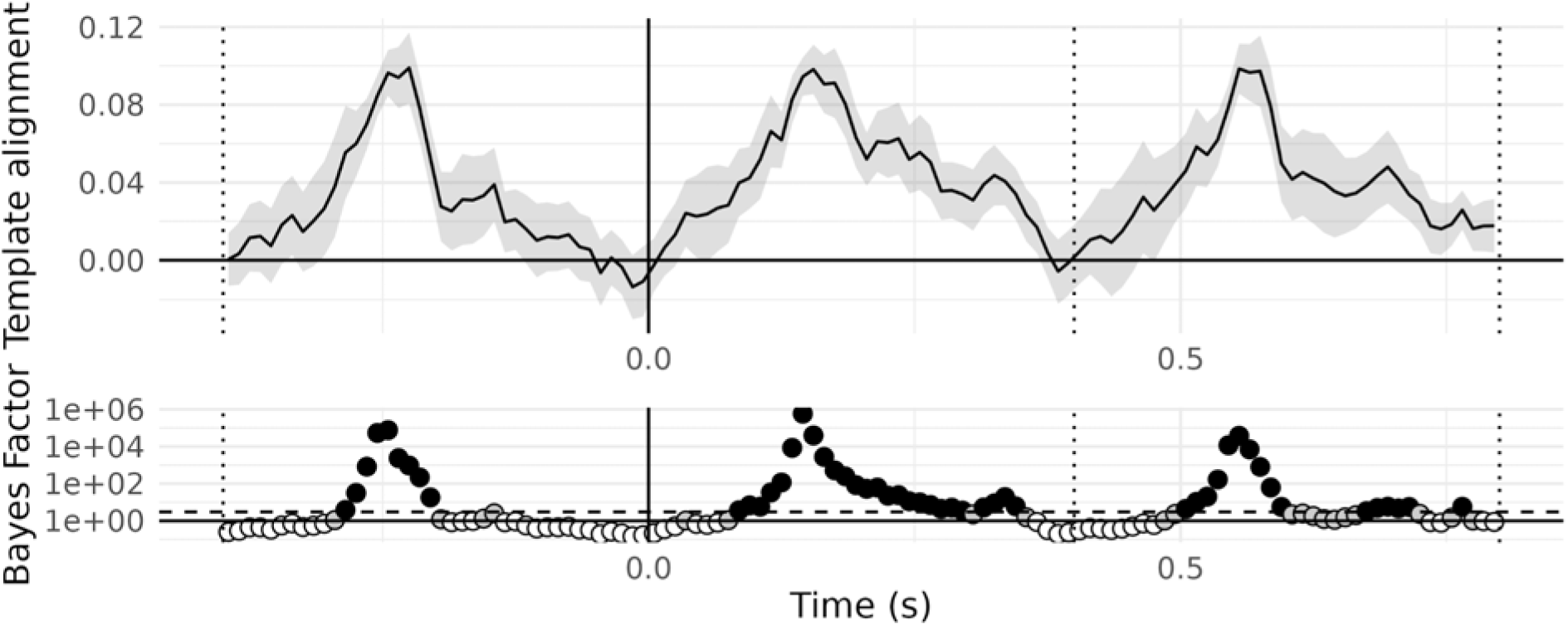
Biased, stimulus-unspecific template alignment. Upper panel: Mean and standard error of (biased) template alignment are plotted for each time point. Lower panel: The corresponding Bayes Factors are plotted, similar to Figure 2D. The graphs illustrate how the template alignment calculation can be biased by effects of th experimental phase, if not corrected via equations 1a and 1b.

### Comparison of different distance metrics

We chose the Mahalanobis distance to quantify the distances between neural representations. Mahalanobis distance has several advantages compared to, for example, the Euclidean distance, as both the variance and the covariance between features is taken into account. This makes particularly sense since both neighboring electrodes share information, as well as neighboring time points. To test the influence of the distance metric on our results, we repeated the main analyses and replaced Mahalanobis distance with Euclidean distance and Correlation distance. For Euclidean distance, each feature (i.e., each time point and channel, across all trials from all conditions) was scaled to follow N(0,1) prior to distance estimation. For Correlation distance, correlations were arctanh-transformed (e.g., eqs. 1-3).

The results indicated evidence for template alignment with both alternative distance metrics, i.e., BF = 15, V = 131, p = 0.004, g = 0.72 [0.20, 1.23] for Euclidean distance, and BF = 8, V = 127, p = 0.007, g = 0.64 [0.13, 1.14] for Correlation distance. Using Euclidean distance a higher category selectivity was observed for TPL stimuli compared to PRE stimuli (BF = 7, V = 130, p = 0.014 (false discovery rate-corrected), g = 0.62 [0.11, 1.11]), but no evidence for the other comparisons or by using Correlation distance (all BF < 3, all p > 0.05). Spontaneous alignment was similarly not found (all BF < 3, all p > 0.05).

### Spatial feature constriction

We repeated all main analyses also by constricting features to only posterior regions of interest, i.e., O1, Oz, O2, POz, P7, P3, Pz, P4, P8, CP5, CP3, CP4, CP6, T7, C3, C4 & T8. We found similar evidence as when using whole-brain features regarding template alignment (BF = 19.4, V = 132, p = 0.003, g = 0.75 [0.22, 1.27]). Category selectivity similarly showed no increase from PRE to POST (p > 0.05), but higher values for TPL compared to PRE and POST phases (all p < 0.001). Similarly, no spontaneous alignment was observed (p > 0.05).

## Supplementary Tables

**Table S1.** Counts of response categories. The responses within the ratings block after PRE and POST phases (Fig. 1) were used to categorize each stimulus for each participant. The counts for each image being categorized in the different groups are illustrated.

| Image | Learned | Re-learned | Spontaneous | Not learned | Not re-learned |
|---|---|---|---|---|---|
| cow_02 | 8 | 8 | 0 | 1 | 0 |
| cow_03 | 7 | 6 | 4 | 0 | 0 |
| croissant_01 | 5 | 11 | 0 | 0 | 1 |
| croissant_03 | 11 | 4 | 2 | 0 | 0 |
| hands_02 | 8 | 9 | 0 | 0 | 0 |
| hands_03 | 5 | 6 | 6 | 0 | 0 |
| human_01 | 2 | 9 | 6 | 0 | 0 |
| human_02 | 5 | 10 | 1 | 1 | 0 |
| penguin_06 | 9 | 6 | 2 | 0 | 0 |
| penguin_07 | 13 | 2 | 0 | 0 | 2 |

**Table S2.** Sources of the stimulus images. Images were taken either from the THINGS database, or otherwise searched for on the internet. The table lists the original sources of the images, license type and author. For two images, the original source is unclear, although they appear to serve widely as stock photos. They are therefore censored in the manuscript, but can be identified over the THINGS database (https://osf.io/jum2f).

| Image | Dataset | THINGS id | Original source | License | Author |
|---|---|---|---|---|---|
| cow_02 | THINGS | calf1_10s | https://pixabay.com/photos/cow-calf-young-animal-beef-790263/ | Pixabay license<sup>1</sup> | Petra “Pezibear” |
| cow_03 | THINGS | calf1_14s | unclear | unclear | - |
| croissant_01 | THINGS | croissant_05s | https://www.freeimageslive.co.uk/free_stock_image/croissant-jpg | CC BY 3.0 | freefoodphotos.com |
| croissant_03 | THINGS | croissant_08s | unclear | unclear | - |
| hands_02 | internet | - | https://www.pexels.com/photo/faceless-artisan-planing-hardwood-plank-5974252/ | Pexels license<sup>2</sup> | Ono Kosuki |
| hands_03 | internet | - | https://www.pexels.com/de-de/foto/holz-industrie-gesichtslos-verwischen-5974289/ | Pexels license | Ono Kosuki |
| human_01 | internet | - | https://www.pexels.com/de-de/foto/madchen-spielt-mit-blasen-1919030/ | Pexels license | Alexander Dummer |
| human_02 | internet | - | https://www.pexels.com/photo/person-swimming-on-body-of-water-863988/ | Pexels license | Guduru Ajay Bhargav |
| penguin_06 | THINGS | penguin_15s | https://pixabay.com/nl/photos/koningspinguïn-wandeling-1154432/ | Pixabay license | Marcel Langthim |
| penguin_07 | THINGS | penguin_20s | https://pixabay.com/photos/penguin-zoo-bird-animal-1598209/ | Pixabay license | Joanna Jankowski |

<sup>1</sup> https://pixabay.com/service/license-summary/
<sup>2</sup> https://www.pexels.com/license/

## Notes

### Competing Interest Statement

The authors have declared no competing interest.

### Summary of Updates

- shortened analysis time window - expanded discussion and interpretation - extended analyses - error corrections

